# A novel head-fixed assay for social touch in mice uncovers aversive responses in two autism models

**DOI:** 10.1101/2023.01.11.523491

**Authors:** Trishala Chari, Ariana Hernandez, Carlos Portera-Cailliau

## Abstract

Social touch, an important aspect of social interaction and communication, is essential to kinship across animal species. How animals experience and respond to social touch has not been thoroughly investigated, in part due to the lack of appropriate assays. Previous studies that examined social touch in freely moving rodents lacked the necessary temporal and spatial control over individual touch interactions. We designed a novel head-fixed assay for social touch in mice, in which the experimenter has complete control to elicit highly stereotyped bouts of social touch between two animals. The user determines the number, duration, context, and type of social touch interactions, while monitoring with high frame rate cameras an array of complex behavioral responses. We focused on social touch to the face because of their high translational relevance to humans. We validated this assay in two different models of autism spectrum disorder (ASD), the *Fmr1* knockout model of Fragile X Syndrome and maternal immune activation mice. We observed increased avoidance, hyperarousal, and more aversive facial expressions to social touch, but not to object touch, in both ASD models compared to controls. Because this new social touch assay for head-fixed mice can be used to record neural activity during repeated bouts of social touch it should be of interest to neuroscientists interested in uncovering the underlying circuits.

## INTRODUCTION

Across animal species and humans, social touch is an important component of social interaction and communication that allows for the development of strong kinship bonds (Adolphs, 2009; Bales et al., 2018; Dunbar, 2010). Touch may be experienced under different contexts, such as between parent and offspring, siblings, friends, or even strangers (Chen & Hong, 2018). Whether animals experience social touch as pleasant or aversive, and the degree to which their behavioral responses differ from those related to touching inanimate objects is not known. Moreover, the neural circuits encoding social touch or how activity within those circuits relates to the behavioral repertoire animals exhibit in response to social touch are not fully understood.

Animal studies have begun to address these questions, especially in rodents, using a variety of behavioral paradigms. Unfortunately, the social touch assays currently available have certain limitations. Those that favor naturalistic interactions in freely moving rodents lack temporal and spatial control over individual touch interactions and typically the data collected reflect a mix of different interactions occurring simultaneously (e.g., anogenital sniffing, whisker-whisker contact, allo-grooming) (Bobrov, Wolfe, Rao, & Brecht, 2014; Jennings et al., 2019; Lenschow & Brecht, 2015; Mosher, Zimmerman, Fuglevand, & Gothard, 2016; Yu et al., 2022). To overcome these problems, we sought to design a novel head-fixed social touch behavioral assay for rodents, in which we could control the duration, number, context, and type of social touch interactions with high precision, while at the same time monitoring an array of complex behavioral responses (facial expressions, pupillary changes, motor avoidance, etc.) using high frame rate cameras. We focused on a single type of social touch interaction (face-to-face), as opposed to the equally prevalent anogenital sniffing interactions in mice (Chen & Hong, 2018; Ebbesen & Froemke, 2022), because we felt it had more translational relevance to humans. We took care to ensure that the experimenter had complete control to directly elicit highly stereotyped bouts of social touch between animals.

To validate our new assay, we used it to identify differences in behavioral responses to social touch in mouse models of Autism Spectrum Disorder (ASD). ASD is a prevalent neurodevelopmental condition characterized by deficits in social interaction, repetitive behaviors, and atypical sensory processing (Robertson & Baron-Cohen, 2017). The change in quality of life in autistic individuals is primarily attributed to social deficits, which can be associated with (or even triggered by) atypical processing of sensory stimuli (Baron-Cohen & Belmonte, 2005; Lee Masson et al., 2019; Thye, Bednarz, Herringshaw, Sartin, & Kana, 2018). Apprehension to social touch in ASD could be caused by tactile hypersensitivity (Cascio et al., 2008; Green & Ben-Sasson, 2010; Green et al., 2015; He et al., 2017), which is a strong predictor of future social deficits (Green, Hernandez, Bowman, Bookheimer, & Dapretto, 2018). Avoidance of social touch by ASD children could prevent them from forming social relationships as adults (Foss-Feig, Heacock, & Cascio, 2012; Thye et al., 2018). In certain rodent models of autism, tactile sensitivity and social interaction deficits also appear to be linked (Orefice et al., 2019; Orefice et al., 2016), and, in some cases, differences in the development of primary somatosensory cortex (S1) are associated with social deficits (Choi et al., 2016; Reed et al., 2020). Thus, further research into social touch in ASD models is warranted.

We tested two distinct mouse models of ASD (the Fmr1 knockout model of Fragile X Syndrome and maternal immune activation mice) in our novel head-fixed social touch assay. We quantified various behavioral responses in the test animal in response to social touch of a stranger mouse. We observed increased avoidance, hyperarousal (pupil dilation), and more aversive facial expressions to social touch (but not to object touch) in both ASD models compared to their healthy controls. Our results suggest that this new social touch assay can parse out maladaptive behavioral responses to social touch in ASD mouse models and might be of use to the larger neuroscience community.

## RESULTS

### A novel behavioral assay for social touch

To investigate how mice respond to social touch, and the circuits involved, one must consider the pros and cons of different behavioral assays. Inspired by prior designs of social touch assays for mice and rats (Bobrov et al., 2014; Jennings et al., 2019), we designed a novel head-fixed behavioral assay in which we can control the frequency and duration of each social touch interaction, the type of touch (whisker-whisker vs. snout-snout), and the context (social vs. object). In this this assay, a head-restrained test mouse that is allowed to run on an air-suspended polystyrene ball is monitored with multiple cameras during repeated presentations of a novel mouse that is also head-fixed and resting on a motorized stage that brings it to predetermined positions at various distances away from the test mouse (see *Methods*, **Fig. 1**). We tested three different positions of the stage to assess corresponding conditions of social touch: 1. No touch, where the test animal can see the novel ‘visitor’ mouse but not touch it; 2. Voluntary social touch where the test mouse can interact with the visitor via its whiskers; 3. Forced social touch, where the visitor mouse is so close to the test mouse that their snouts are in direct physical contact.

**Fig. 1:**
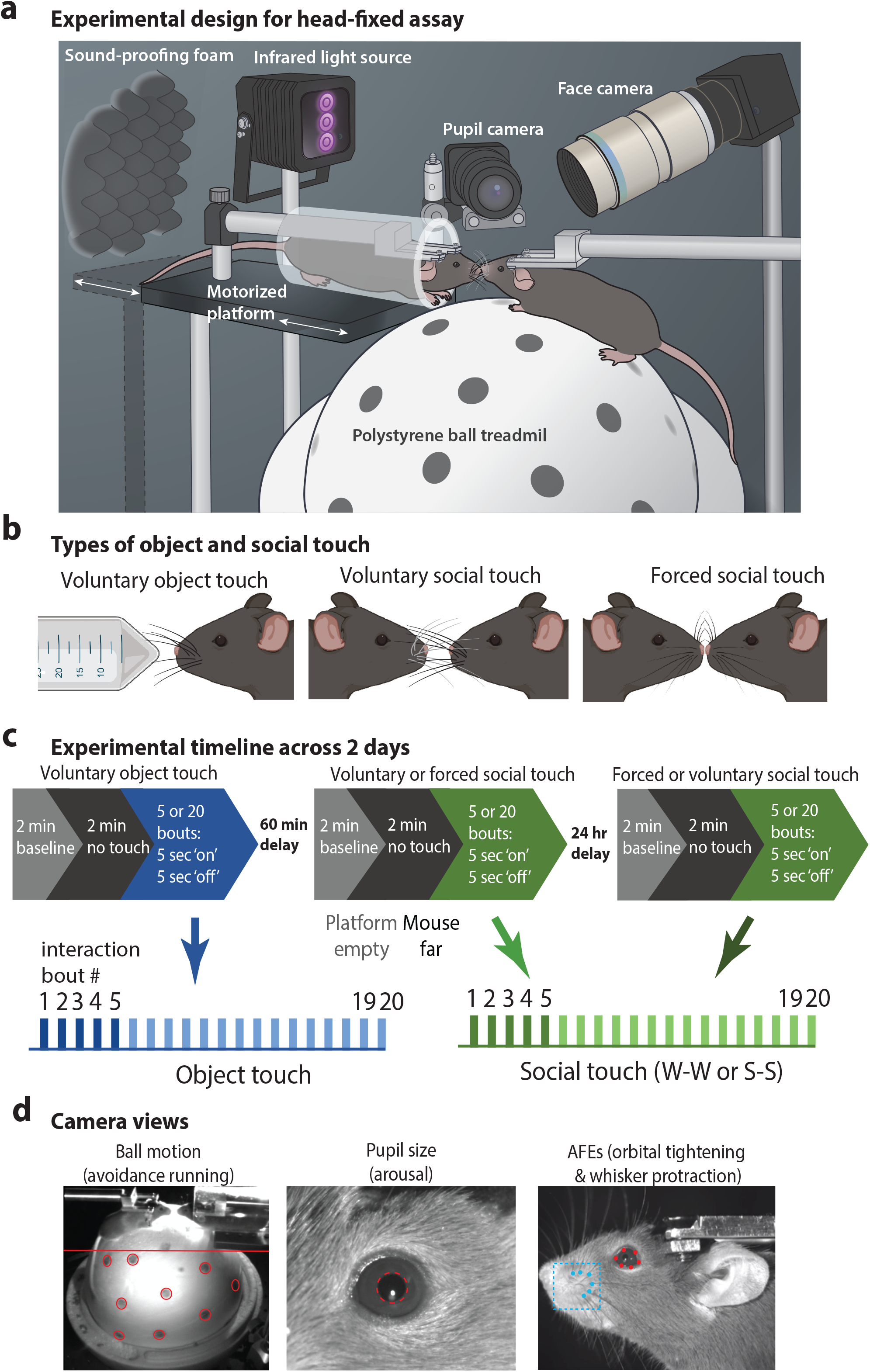
Setup for social touch behavioral assaya. **a**.Overview of head-fixed setup for the social touch behavioral assay. A head-fixed test mouse can run on an air-suspended polysterene ball while interacting with a stranger mouse restrained in a plexiglass tube secured to a motorized platform. The system is fully automated to move the stranger mouse to different distances away from the test mouse. Two cameras focus on the face and the eye/pupil, respectively, while a third camera that tracks the mouse and ball motion is overhead (not shown). An infrared light source provides optimal light for tracking behavioral responses. Acoustic foam is used for sound insulation. **b**. We tested three types of touch: voluntary object (whisker-object), voluntary social (whisker-whisker), and forced social (snout-snout). **c**. Duration of baseline (platform empty without object/mouse), no touch, and object social touch, as well as the number of stimulations and delay between each type of touch condition. **d**. Camera views for tracking ball motion, pupil size and AFEs.

By using high frame rate cameras to record the test animal’s face and eyes, as well as ball motion, we can quantify different aspects of facial expressions (e.g., whisker movements, mouth opening, ear movements, eye size changes) and changes in pupil diameter or saccades, as well as locomotion (**Fig. 1d**). Because we are interested in autism, we focused on behaviors that might indicate that the mouse experienced social touch as an aversive stimulus, by exhibiting avoidance, defensive behaviors, emotional facial expressions, or hyperarousal. Indeed, these behavioral responses are observed in ASD individuals responding to social or affective touch and in mouse models of ASD responding to passive non-social touch (Bales et al., 2018; Cascio et al., 2008; He et al., 2017; Klusek, Martin, & Losh, 2013; Mammen et al., 2015; Thye et al., 2018; Zampella, Bennetto, & Herrington, 2020). Our assay also examines social touch that is potentially unpleasant, rather than allo-grooming, by including forced snout-snout interactions. This allowed us to explore how ASD mouse models might respond to social touch across different contexts and how the tactile system engages with these stimuli behaviorally. However, this social touch assay can be easily modified to change the presentation parameters, the types of visitor and test mice, in order to explore a myriad of interesting questions about social touch in rodents. We also designed the assay to be compatible with silicon probes or calcium imaging recordings of neural activity, to elucidate circuits that are activated by social touch, as well as those that mediate behavioral responses to social touch.

To demonstrate the utility of this novel social touch assay, we compared the behavioral responses of control wild-type mice to those of two mouse models of ASD. The first was the *Fmr1* knockout (*Fmr1*^*-/-*^) mouse model of Fragile X Syndrome (FXS) (Dutch-Belgian Fragile X consortium), the leading single gene cause of intellectual disability and autism. The other was the Poly(I:C) maternal immune activation (MIA) model (**Fig. 1 Suppl. Fig. 1**), which is widely used as a model of an environmental cause of autism (Choi et al., 2016; Estes & McAllister, 2016; Kentner et al., 2019). Here, we present results of our observations related to four major behavioral responses: 1. Avoidance running; 2. Pupil dilation; 3. Whisker protraction; and 4. Orbital tightening (squinting). Overall, we hypothesized that, compared to their respective controls, *Fmr1*^*-/-*^ and MIA mice would show increased avoidance, hyperarousal, and more aversive facial expressions (AFEs) to social touch, but no differences for object touch. Furthermore, we expected that forced social touch (snout-snout) would be more aversive than voluntary social interactions (whisker-whisker) for ASD mice.

### Greater avoidance running in Fmr1^-/-^ and MIA mice during social touch but not object touch

Sensory hypersensitivity is very prevalent in ASD and is thought to contribute to maladaptive avoidance responses, such as tactile defensiveness and social avoidance (Baranek, Foster, & Berkson, 1997; Robertson & Baron-Cohen, 2017). Most children with FXS experience sensory over-reactivity, often leading to tactile defensiveness and gaze aversion (Rais, Binder, Razak, & Ethell, 2018; Sinclair, Oranje, Razak, Siegel, & Schmid, 2017). However, avoidance to social touch per se has never been investigated in animal models of ASD or FXS. Previously, we demonstrated that *Fmr1*^*-/-*^ mice exhibit tactile defensiveness to repetitive whisker stimulation, which manifested as avoidance running(He et al., 2017). To investigate whether social touch leads to avoidance, we quantified running direction of the test mouse relative to the novel object or stranger mouse (**Fig. 2a, Fig. 2 Suppl. Fig. 1a**). If the test animal was moving backward (either left or right), we categorized this as avoidance, in contrast to running forward, which was considered an adaptive response (seeking social interaction). We initially calculated the total time the mouse spent in locomotion, regardless of direction, to determine if group differences in running might skew the proportion of avoidance running. Although there are reports of hyperactivity in *Fmr1*^*-/-*^ mice (“Fmr1 knockout mice: a model to study fragile X mental retardation. The Dutch-Belgian Fragile X Consortium,” 1994; Sullivan et al., 2006), we have not found differences in total locomotion between adult *Fmr1*^*-/-*^ and WT mice either in response to whisker stimulation or while performing a visual discrimination task (Goel et al., 2018; He et al., 2017). In the social touch assay, we observed that mice of all groups spent more time running when they transitioned from the baseline period (before touch) to the period of social touch (**Fig. 2 Suppl. Fig. 1b**). There were no differences in running speed during object or social touch (**Fig. 2b**), or in locomotion between *Fmr1*^*-/-*^ or MIA mice and their respective controls (**Fig. 2 Suppl. Fig. 1b**). Therefore, locomotion is similar between the two ASD models and their controls both before and during social touch, so any differences in avoidance running could not be explained by hyperactivity.

**Fig. 2:**
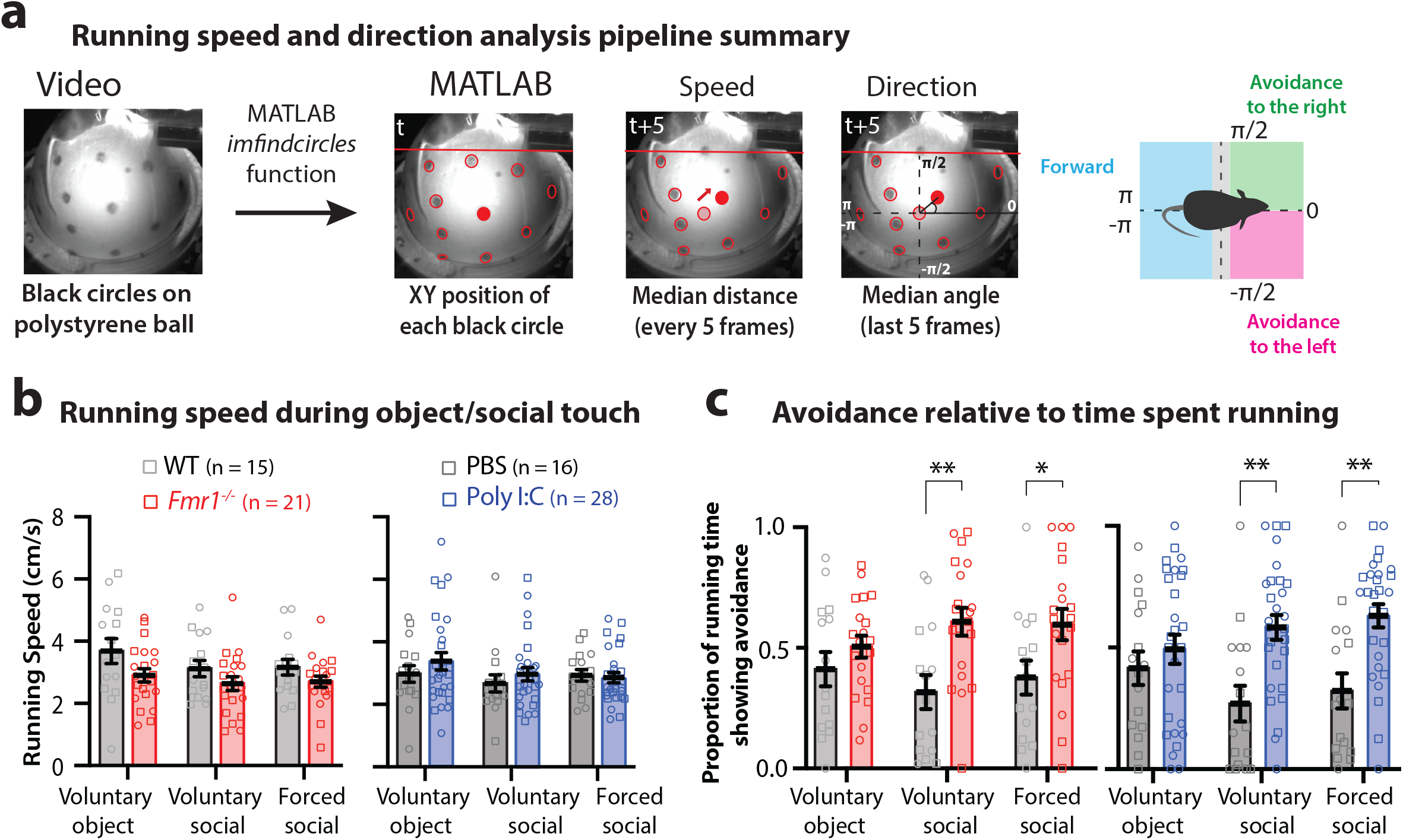
Mouse models of autism show avoidance to social touch from a stranger mouse. **a**. Summary of analysis for running speed and direction (see also Suppl Fig. 1). **b**. Running speeds during all types of touch do no differ between ASD mice and control animals. **c**. Running avoidance (backwards to left or right) is higher in *Fmr1*^*-/-*^ and Poly(I:C) MIA mice compared to controls during voluntary and forced social touch but not object touch. Squares=males, circles females. **p<0.01, *p<0.05, two-way ANOVA with Bonferroni’s.

When we compared the time spent showing avoidance running as a proportion of total locomotion, we found that both *Fmr1*^*-/-*^ and MIA mice displayed higher avoidance during voluntary and forced social touch compared to controls, but not during voluntary object touch (**Fig. 2**; WT vs. *Fmr1*^*-/-*^: vol. object p>0.05, vol. social p=0.0045, forc. social p=0.0467; PBS vs. MIA: vol. object p>0.05, vol. social p=0.0017, forc. social p=0.0021). ASD mice, but not controls, also showed more avoidance running during social touch than just before touch, but not during object touch (**Fig. 2 Suppl. Fig. 1c**; WT vs. *Fmr1*^*-/-*^: vol. social p=0.0042, forc. social p<0.001; PBS vs. MIA: vol. social p<0.001, forc. social p=0.0029).

Because there are important sex differences in both the prevalence and symptoms of ASD (Bartholomay, Lee, Bruno, Lightbody, & Reiss, 2019; Werling & Geschwind, 2013), we compared avoidance between male and female mice in each group but did not find any significant differences even though avoidance in WT and PBS females appears a little higher (**Fig. 2 Suppl. Fig. 2a**, p>0.05). We also looked at differences between litters in each group but did not see any obvious differences either, although the sample size per litter was small. We only observed one litter in WT (L3) and one in MIA (L3) group with similar levels of avoidance as their comparison groups during object and social touch (**Fig. 2 Suppl. Fig. 2b**; shaded symbols). Interestingly, litter L3 of the MIA group came from a Poly(I:C) injected dam with the lowest IL-6 levels (**Fig. 1 Suppl. Fig. 1a**), and IL-6 levels have been shown to correlate with deficits in this model(Garay, Hsiao, Patterson, & McAllister, 2013; Kentner et al., 2019).

### Pupil dilation with social touch lasts longer in Fmr1^-/-^ and MIA mice

Autonomic hyperarousal, including elevated heart rate and pupil dilation, is observed in autistic individuals during tactile stimulation or affective touch, and is used as an indicator of tactile hypersensitivity (Fukuyama, Kumagaya, Asada, Ayaya, & Kato, 2017; Heilman, Harden, Zageris, Berry-Kravis, & Porges, 2011; McGlone, Wessberg, & Olausson, 2014). We measured changes in pupil size as a proxy for arousal in response to social touch (**Fig. 3a, Fig. 3 Suppl. Fig. 1a, Movie 4**). We first compared pupil size as a mean of the first 5 presentations and found no differences between WT and *Fmr1*^*-/-*^ or between PBS and MIA mice, regardless of condition (object or social touch) (**Fig. 3 Suppl. Fig. 1b**, p>0.05). Next, because pupils can dilate or constrict over short time scales, we compared pupil size at individual presentations of object or social touch (Joshi & Gold, 2020; Vinck, Batista-Brito, Knoblich, & Cardin, 2015). We found that in all groups pupils significantly dilated to a similar extent after the first object/mouse presentation (**Fig. 3b)**. Interestingly, after repeated presentations of voluntary object touch, pupil size returned to baseline in all groups, presumably as a form of adaptation to a non-threatening situation (**Fig. 3b; Fig. 3 Suppl. Fig. 1b**). In contrast, pupils remained dilated for a longer period in *Fmr1*^*-/-*^ mice with both voluntary social touch and forced social touch, and in MIA mice with forced social touch, whereas they constricted to baseline in controls. The difference was most pronounced with forced social touch, where pupils were significantly larger in ASD mice than in their controls on the 5^th^ presentation (**Fig. 3b**; pupil size: WT vs. *Fmr1*^*-/-*^ p<0.001, PBS vs. MIA p=0.0367). In a subset of mice that we tested up to 20 presentations of object and social touch, we found that pupil size in *Fmr1*^*-/-*^ and MIA mice eventually returned to baseline (**Fig. 3 Suppl. Fig. 1b**). We did not find any sex or litter differences in pupil size before or after social touch (**Fig. 3 Suppl. Fig. 2**). Altogether, these findings indicate that both *Fmr1*^*-/-*^ and MIA mice display more avoidance and hyperarousal than their controls to social touch, but not to object touch.

**Fig. 3:**
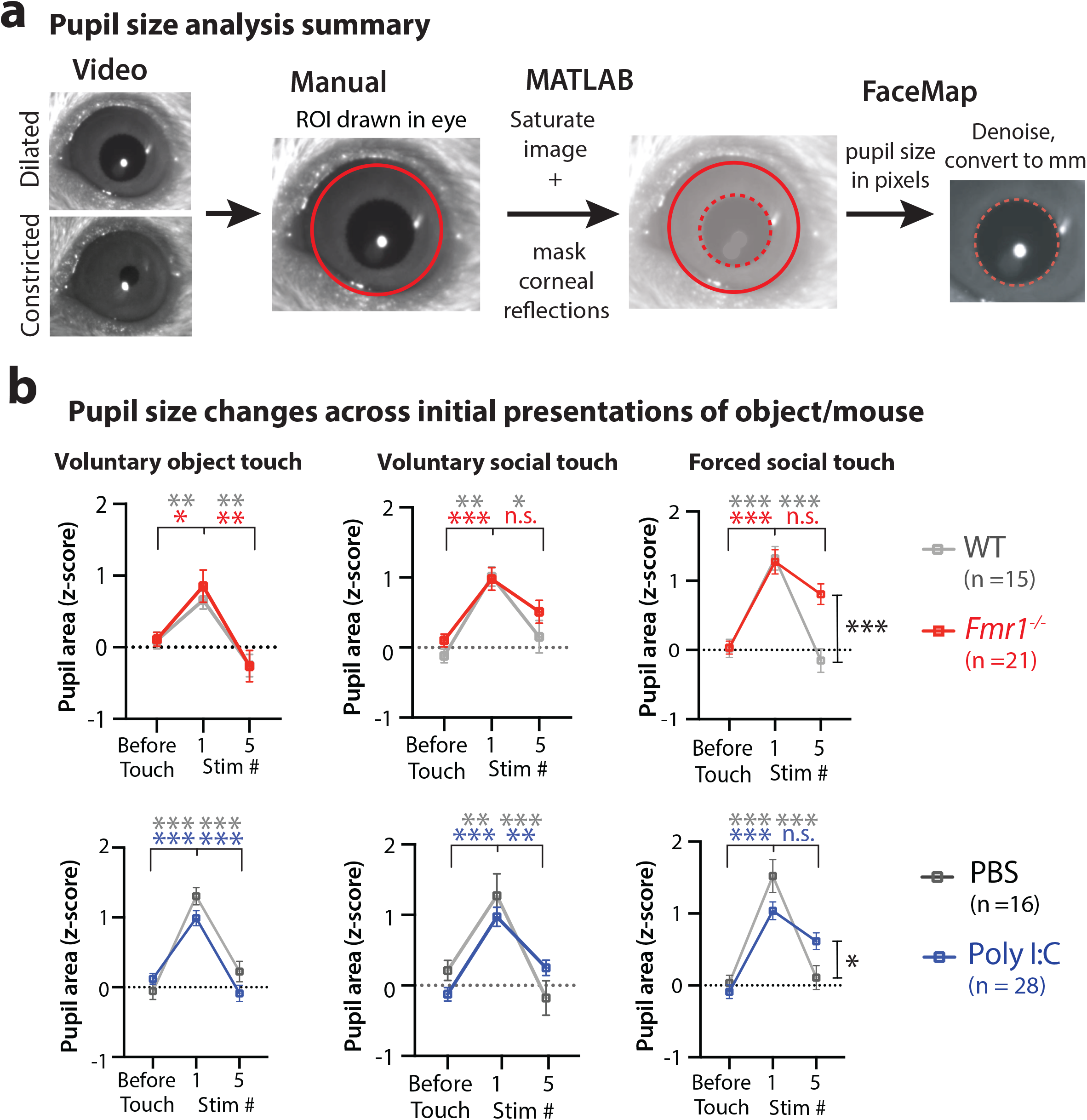
Pupil dilation is prolonged during social touch in ASD mice. **a**. Summary of pupil size analysis using Facemap (see Suppl. Fig. 1). A region of interest (ROI) is drawn in the Facemap graphical user interface in Python (red circle). Facemap detects the pupil within the ROI (red dashed circle) and generates pupil area in pixels, which is converted to mm in MATLAB. **b**. Pupil size does not decrease to baseline levels (before touch) in *Fmr1*^*-/-*^ and MIA mice by the 5th stimulation of voluntary and forced social touch. In all control groups pupil size decreases to before touch at the 5th stimulation for all touch conditions. ***p<0.001, ** p<0.01, *p<0.05, two-way ANOVA with Bonferroni’s for pupil area before touch vs. 1st and 5th stimulation.

### Aversive facial expressions (grimace) are more pronounced in Fmr1^-/-^ and MIA mice during forced social touch

In humans, facial expressions are considered good indicators of emotional state (Anderson & Adolphs, 2014). Autistic individuals will grimace or wince to aversive sensory stimuli and will avert their gaze during social interactions (Foss-Feig et al., 2012; Kliemann, Dziobek, Hatri, Steimke, & Heekeren, 2010; Schmitt, Cook, Sweeney, & Mosconi, 2014). Facial grimacing in the form of orbital tightening, changes in whiskers, nose bulging is also observed in rodents while experiencing pain (Langford et al., 2010), but less is known about facial expressions associated with sensory hypersensitivity or unwanted social interactions. We posited that if ASD mice consider social touch as aversive, they would manifest aversive facial expressions (AFEs). We focused on two facial features, whisker movement and orbital tightening (**Movie 5**), because they were easily detectable by cameras in our set-up and because analysis could be automated using *DeepLabCut* (Langford et al., 2010; Mathis et al., 2018). For whisker movement, we quantified bouts of sustained whisker protraction (**Movie 5**), which is often seen in mice experiencing pain (Langford et al., 2010), in mice subjected to tail shocks, and during aggression or immediate facial contact (Defensor, Corley, Blanchard, & Blanchard, 2012; Dolensek, Gehrlach, Klein, & Gogolla, 2020; Ebbesen & Froemke, 2021; Wolfe, Mende, & Brecht, 2011). Prolonged whisker protraction is different from active whisking, which is an adaptive behavior in rodents as they explore their environment, both in terms of the speed and the direction of whisker movement. During active whisking, follicles are displaced forwards and backwards rapidly and rhythmically (8-12 Hz, for bouts lasting 1-2 seconds) (Bush, Solla, & Hartmann, 2016). In contrast, during aversive whisker protraction, the animal’s whiskers are maintained in a fixed, forward position, for bouts lasting several seconds.

We could distinguish between these two types of whisker movement using *DeepLabCut* and *Facemap* (**Fig. 4a, Fig. 4 Supp Fig. 1a-b**; see Methods). ASD mice and their controls all showed more active whisking when presented with novel mice compared to baseline (**Fig. 4 Suppl. Fig, 1c**), but we did not find any significant differences in time spent actively whisking between groups or between voluntary or forced social touch. However, we found that *Fmr1*^*-/-*^ and MIA mice spent significantly more time than their controls displaying aversive whisker protraction during forced social touch (**Fig. 4b**, WT vs. *Fmr1*^*-/-*^ p<0.001, PBS vs. MIA p=0.0063). In contrast, we saw no group differences in whisker protraction during object touch or voluntary social touch. There were no significant sex or litter differences in whisker protraction in any group across all touch conditions (**Fig. 4 Suppl. Fig. 2**, p>0.05).

**Fig. 4:**
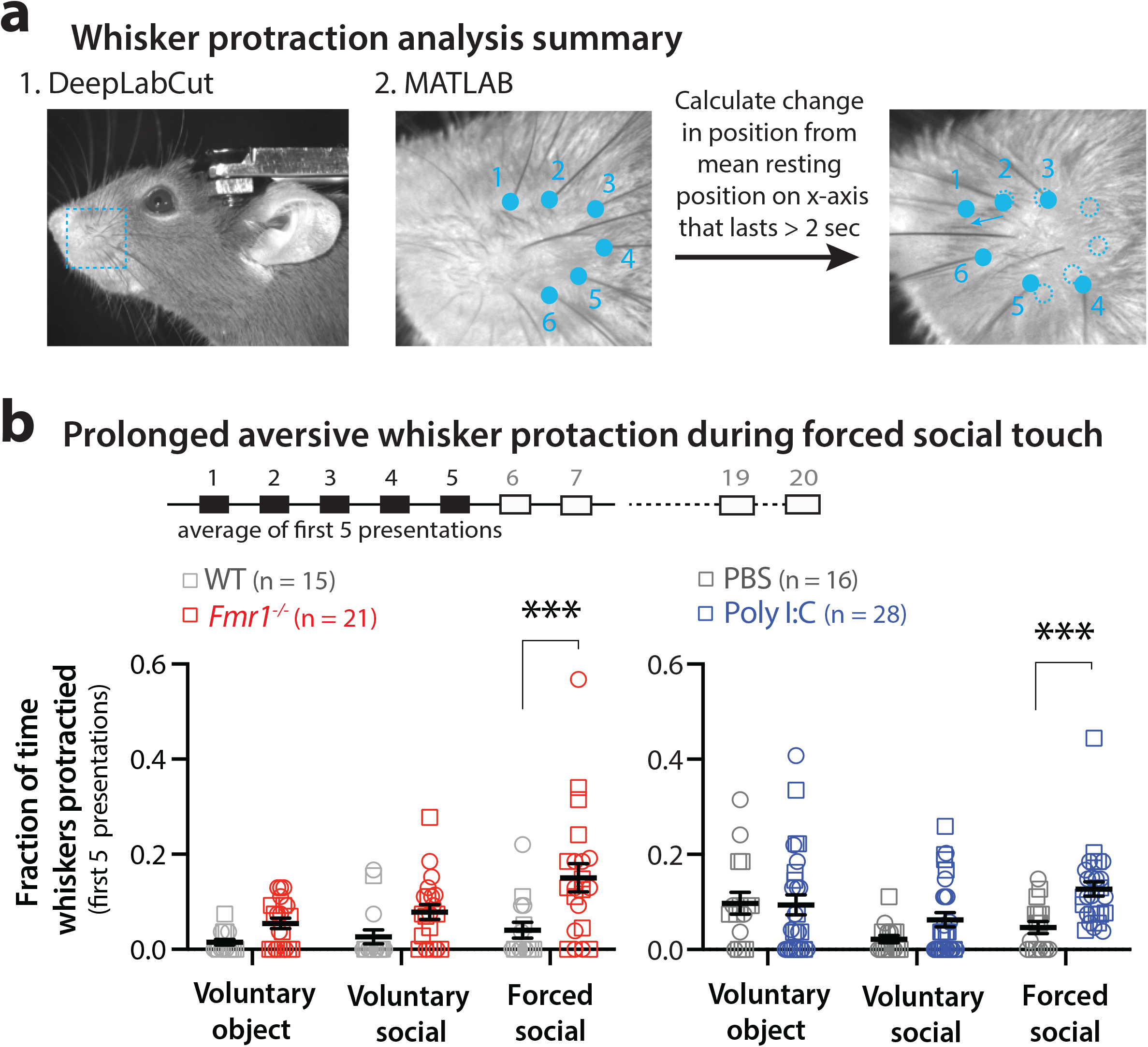
Prolonged whisker protraction during forced social touch in ASD mice. **a**. Summary of analysis for calculating prolonged whisker protraction. Whisker protraction is determined by training a deep neural network (DNN) in DeepLabCut to detect 6 whisker follicles on videos of the mouse’s face (see Suppl. Fig. 1). **b**. The fraction of time *Fmr1*^*-/-*^ and MIA mice exhibited prolonged whisker protraction was higher during forced social touch than their controls but not significantly for voluntary object and social touch. Squares=males, circles=females.***p < 0.001 for two-way ANOVA with Bonferroni’s.

Next, we determined whether mice show orbital tightening during social touch by estimating the area of the eye during the first 5 presentations of social touch. Again, we used *DeepLabCut* to train a neuronal network to estimate the area of the eye (**Fig. 5a, Fig. 5 Suppl. Fig. 1a)**. We omitted frames in which the mouse would blink or groom the face with its forepaws (thereby obscuring the eye) by visual inspection (see *Methods*). We found that orbital area significantly decreased (i.e., more orbital tightening) in *Fmr1*^*-/-*^ and MIA mice during forced social touch, but not during voluntary object or voluntary social touch, and not in the WT or PBS controls (**Fig. 5b**, WT vs. *Fmr1*^*-/-*^ p=0.0418, PBS vs. MIA p<0.001). The orbital area for both *Fmr1*^*-/-*^ and MIA mice was also significantly smaller than in their controls during forced social touch compared to just before touch (**Fig. 5c**, WT vs. *Fmr1*^*-/-*^ p=0.0193, PBS vs. MIA p=0.0188). However, this was not so for voluntary social touch. No differences in orbital tightening were found between sex or between litters in any group or across all touch conditions (**Fig. 5, Suppl. Fig. 1b-c**, p>0.05). These findings suggest that AFEs like sustained whisker protraction and forceful eye closure are uniquely triggered by forced social interactions in ASD mouse models.

**Fig. 5:**
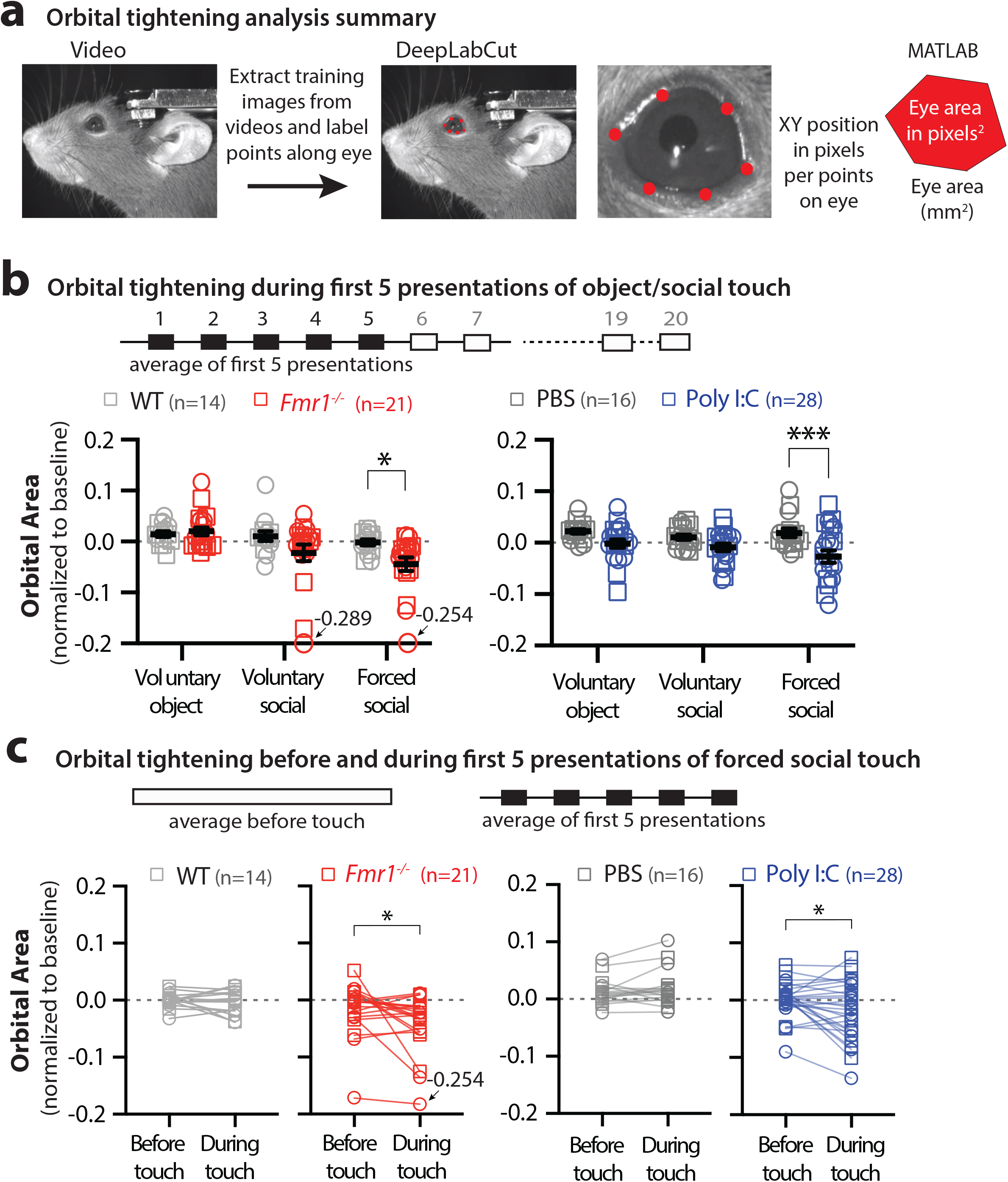
Orbital tightening during forced social touch in ASD mice. **a**. Summary of analysis for calculating orbital tightening. Whisker protraction is determined by training a DNN in DeepLabCut to detect 6 points along the eye on videos of the mouse’s face (see Suppl. Fig. 1). **b**. Orbital eye during touch is normalized to area at baseline (no object or mouse in behavior rig). Orbital area is significantly lower (greater orbital tightening) during forced social touch in *Fmr1*^*-/-*^ and MIA mice compared to controls. **c**. Orbital area is significantly lower during touch compared to the time period before touch in both *Fmr1*^*-/-*^ and MIA mice. Squares=males, circles=females. ***p< 0.001, **p<0.01, *p<0.05, two-way ANOVA with Bonferroni’s. ROUT outlier test removed one animal from WT group.

## DISCUSSION

The main goal of this study was to implement a new behavioral paradigm to investigate social touch behaviors in rodents and the underlying circuits involved. Our findings can be summarized as follows: 1. Our new social touch assay can reliably distinguish behavioral responses of mice to social touch from their responses to inanimate objects; 2. Relative to typically developing control mice, both *Fmr1*^*-/-*^ and MIA mice show increased avoidance running to both voluntary and forced social touch, but not to object touch; 3. Hyperarousal (as measured by pupil dilation) to social touch lasts longer in ASD mice compared to their controls; and 4. AFEs to social touch are more pronounced in ASD mice than in controls, especially during forced social touch.

A few prior studies investigated social touch in freely moving rodents (Bobrov et al., 2014; Jennings et al., 2019; Yu et al., 2022). Despite their ingenuity, the assays relied on at least one animal initiating social touch, and they could not focus on one particular aspect of social touch (e.g., face-to-face contact) amongst the broad and complex behavioral repertoire (e.g., ano-genital sniffing, allo-grooming). Moreover, while somewhat more naturalistic in their design, those assays were limited by the fact that individual social touch interactions varied in duration and frequency, and the timing could not be anticipated by the experimenters. We purposely designed a new assay for head-fixed rodents so the experimenter could control all aspects of the social touch interaction, from the duration and number of interactions to the context of the interaction (voluntary vs. forced, object vs. social). This allowed us to monitor various behavioral responses of the animal to social touch, including body movements that indicated avoidance, facial expressions suggestive of aversion, and dilated pupils reflecting hyperarousal and anxiety. We could even track rapid saccades consistent with gaze avoidance (not shown). Importantly, our assay can easily be combined with 2-photon calcium imaging and silicon probes to record neural activity during social interactions. Because the social touch presentations are highly stereotyped across large numbers of trials, the data from neural recordings would be highly reproducible, and one could quantify the degree to which neurons adapt their responses to repeated presentations. Thus, our assay should be of help to neuroscientists interested in investigating social behaviors in rodents and the circuits involved.

To validate this assay we probed social touch within a disease context in which social deficits are observed, by examining two mouse models of ASD. A major gap in our understanding of ASD, particularly when using mouse models, concerns the relationship between tactile hypersensitivity and social deficits (Lee Masson et al., 2019; Suvilehto, Glerean, Dunbar, Hari, & Nummenmaa, 2015; Thye et al., 2018). Therefore, we used our social touch assay to characterize three maladaptive behavioral responses to social touch in well-established mouse models of ASD: avoidance running, hyperarousal, and AFEs.

We previously reported avoidance to repetitive whisker stimulation in *Fmr1*^*-/-*^ mice (He et al., 2017). However, avoidance to social touch was not simply a manifestation of generalized sensory hypersensitivity (tactile defensiveness) because it occurred only in the context of social touch and not object touch (**Fig. 2c**). Similar sensory avoidance is also observed in humans with ASD and FXS (Green & Ben-Sasson, 2010; Mammen et al., 2015; Rais et al., 2018). Escape or avoidance has been described in mice responding to threatening stimuli, or those causing discomfort, anxiety or pain (Gehrlach et al., 2019; Huang et al., 2019; La-Vu, Tobias, Schuette, & Adhikari, 2020; Yilmaz & Meister, 2013). It will be important to determine whether other avoidance behaviors, such as defensive grooming or gaze avoidance, can also be detected using our social touch assay (Kleberg et al., 2017; Stuart, Whitehouse, Palermo, Bothe, & Badcock, 2022).

During social touch, ASD model mice exhibited sustained pupil dilation, particularly for forced touch, whereas control mice only showed a transient dilation at the onset of social touch (**Fig. 3b**). Pupil size is commonly used as an indicator of arousal levels and autistic/FXS individuals show deficits in autonomic responses and hyperarousal (Joshi & Gold, 2020; Klusek et al., 2013; Kushki, Brian, Dupuis, & Anagnostou, 2014; Vinck et al., 2015). Interestingly, some autistic individuals who do not exhibit hyperarousal fail to show sensory hypersensitivity (Rogers & Ozonoff, 2005), suggesting that the two phenomena are strongly correlated. However, aside from pupil size, hyperarousal can manifest with other autonomic responses, such as increased heart rate, perspiration, or changes in breathing(Heilman et al., 2011; Kushki et al., 2014), which could also be monitored in our assay.

Finally, we observed that ASD mice exhibit more pronounced AFEs (whisker protraction and orbital tightening), but only during forced social touch. Our findings align well with previous findings concerning facial grimacing in mice (Defensor et al., 2012; Dolensek et al., 2020; Ebbesen & Froemke, 2021; Langford et al., 2010). Autistic people are often unable to recognize or imitate facial expressions of others, which complicates their interactions in social settings (Drimalla, Baskow, Behnia, Roepke, & Dziobek, 2021). We did not monitor the facial expressions of the stranger mice, but the camera setup could be modified to track this too.

We did not find significant sex differences in our assay. This was surprising given that the prevalence of ASD and the range of phenotypic behaviors are different in males and females (Werling & Geschwind, 2013). It is possible that sex differences were not apparent in our head-fixed social touch paradigm because mice could not freely choose to engage social investigation is eliminated. However, our assay could easily be modified to allow the test mouse to exert control of the motorized stage. Interestingly, females showed similar changes in pupil dilation, suggesting the autonomic system is not differentially affected by social touch between males and females. Females also showed similar amounts of whisker protraction as males, suggesting that either females display aggression-like behaviors during touch or that the protraction observed during touch is associated with aversion or defensiveness more so than social aggression.

We recognize that our social touch assay has some limitations. Compared to assays for freely moving mice, our assay is less naturalistic. In spontaneous social interactions, mice are free to decide when to approach another animal. They may choose to approach other mice from the rear, as opposed to face-to-face. Our head-fixed assay also prevents the mice from engaging in other socially relevant behaviors that involve touch, such as allo-grooming, or fighting. Head-fixation also prevents head movements that may be important for mice to engage in social touch.

In summary, our novel head-fixed paradigm revealed that ASD mouse models manifest multiple maladaptive responses to social touch and that these behavioral align well with symptoms and atypical behaviors observed in autistic humans. The fact that two rather distinct ASD models exhibited very similar behavioral phenotypes in avoidance, arousal and facial expressions suggests that our assay may uncover remarkable phenotypic convergence in social touch deficits in other ASD models. Future studies could also explore social touch in other contexts, such as mother-to-pup interactions, or age dependent differences. Finally, one could utilize this assay in combination with in vivo 2-photon calcium imaging or silicon probes to explore changes in neural activity in relevant brain circuits in mouse models of neurodevelopmental conditions.

## MATERIALS AND METHODS

### KEY RESOURCES TABLE

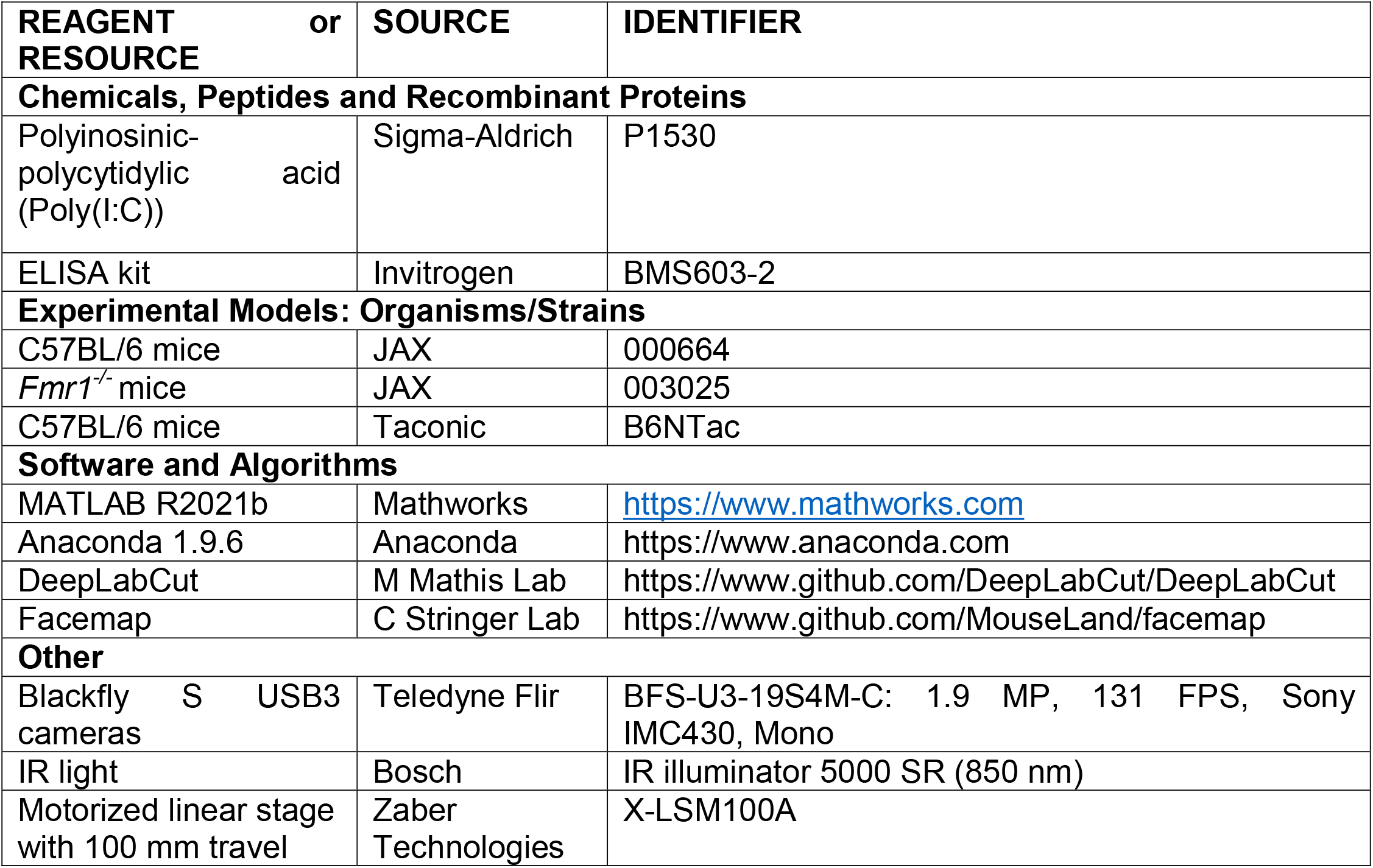

### RESOURCE AVAILABILITY

#### Lead contact

Further information and requests for resources should be directed to and will be fulfilled by the lead contact, Dr. Carlos Portera-Cailliau.

#### Materials availability

This study did not generate new unique reports.

#### Data and code availability

Data will be made available from lead contact, Dr. Carlos Portera-Cailliau upon request so that data can be provided in format most suitable to the requester.

### EXPERIMENTAL MODEL AND SUBJECT DETAILS

#### Animals

Male and female C5BL/6 mice at postnatal day 60-90 on the day of behavioral testing were used for behavioral experiments and were derived from the following mouse lines based on prior publications: wildtype B6J (JAX line 000664), *Fmr1*^*-/-*^ (JAX line 003025), and wild-type (WT) B6NTac (Taconic line)(Choi et al., 2016; “Fmr1 knockout mice: a model to study fragile X mental retardation. The Dutch-Belgian Fragile X Consortium,” 1994; He et al., 2017; Kentner et al., 2019; Reed et al., 2020). The group/genotypes used for behavioral testing are as follows: WT and *Fmr1*^*-/-*^ mice (JAX line) and PBS and maternal immune activation (MIA) mice (Taconic line). Mice were group-housed with access to food and water *ad libitum* under a 12 hour light cycle (12 hours light/12 hours dark) in controlled temperature conditions. All experiments were done in the light cycle and followed the U.S. National Institutes of Health guidelines for animal research under an animal use protocol ARC #2007-035 approved by the Chancellor’s Animal Research Committee and Office for Animal Research Oversight at the University of California, Los Angeles.

#### Maternal immune activation (MIA)

We followed established protocols(Estes & McAllister, 2016; Kentner et al., 2019). Wildtype B6NTac pregnant dams were injected intraperitoneally with polyinosonic:polycytidylic acdi (Poly(I:C)) for MIA or with phosphate buffered saline (PBS; control) at embryonic day 12.5 (E12.5). A small blood sample of the dams was collected from the submandibular vein 2.5 h after injection and centrifuged to isolate serum. Serum was run through an interleukin-6 (IL-6) enzyme-linked immunoabsorbent assay (ELISA) kit (Invitrogen). Successful maternal immune activation in offspring of Poly(I:C) injected dams was confirmed by demonstrating significantly elevated levels of interleukin 6 (IL-6) in the dams compared to PBS-injected dams (Garay et al., 2013) (**Fig. 1 Suppl. Fig. 1a**).

#### Characterization of MIA model

To first characterize that the MIA model we tested whether progeny of Poly(I:C)-injected dams exhibit behavioral deficits previously observed in this model (Choi et al., 2016; Estes & McAllister, 2016; Kentner et al., 2019; Shin Yim et al., 2017). We tested their offspring (male and female) in a battery of behavioral assays (however, these offspring were not tested in the social touch assay). The MIA offspring were tested for the presence of ultrasonic vocalizations at P7-9, in the 3-chamber social interaction assay (which quantifies preference to a novel mouse over an animate novel object) at P60-90, and in the marble burying assay (a measure of repetitive behaviors in rodents) also at P60-90 (Choi et al., 2016; Deacon, 2006; Shin Yim et al., 2017; Yang, Silverman, & Crawley, 2011). MIA mice exhibited reduced pup USV calls, reduced social preference, and increased marble burying compared to PBS offspring. PBS offspring showed no significant behavioral deficits in the 3-chamber and marble burying assays, and their ultrasonic vocalizations as pups, were in the typical range (**Fig. 1 Suppl. Fig. 1b**).

#### Surgical implantation of head bars

Adult mice were anesthetized with isoflurane (5% induction, 1.5-2% maintenance via nose cone v/v) and secured on a stereotaxic frame (Kopf) via metal ear bars. A 1 cm long midline skin incision was made above the skull under sterile conditions. A titanium U-shaped head bar (3.15 mm wide x 10 mm long) was placed on the skull just caudal to Lambda and permanently glued with dental cement. This bar was later used to secure the animal to a post for the head-fixed social touch behavioral assay. This surgery lasted ∼15-20 min and mice fully recovered within 30 min after surgery and returned to group-housed cages.

### BEHAVIORAL ASSAY DETAILS

#### Social touch assay in head-restrained mice

Following head bar implantation, mice were habituated to head restraint and to running on an air-suspended 200 mm polystyrene ball, as well as to the movement of a motorized stage that was used for repeated presentations of an inanimate object or a stranger mouse. The stage was controlled through MATLAB (Mathworks) in a custom-built, sound-attenuated behavioral rig (93 cm x 93 cm x 57 cm) that was dimly illuminated by two infrared lights (Bosch, 850 nm) (**Fig. 1a**). For habituation, test mice were placed on the ball for 20 min each day for 7-9 consecutive days before testing. In parallel, ‘visitor’ mice (stranger to the test mouse) were habituated to head-restraint in a plexiglass tube (diameter: 4 cm) secured to a motorized stage consisting of an aluminum bread board (15 × 7.6 × 1 cm) attached to a translational motor (Zaber Technologies, X-LSM100A). The stage translated at a constant speed of 1.65 cm/s. The neutral starting position was 6 cm away from the test mouse.

Following habituation, test mice were subjected to both voluntary and forced interactions with a visitor mouse or a novel inanimate object over the course of 2 d (**Fig. 1b**). Voluntary interactions meant that the test mouse was within whisker contact of the novel object or mouse, while in forced interactions the stage stopped at a position closer to the test mouse such that the tip of the object or snout of the visitor mouse was in direct contact with the snout of the test mouse. These positions were calibrated before each experiment. On day 1, test mice were placed on the ball and recorded for a 2 min baseline period. Next, we inserted a novel plastic object (a 50 mL Falcon tube) into the plexiglass tube on the motorized stage. For this control interaction the test mouse first experienced a 2 min period of no touch but was able to visualize the object in the neutral position (6 cm away). Next, the motorized stage moved the object to within whisker reach of the test mouse for a total of 20 such presentations of voluntary object touch (**Movie 2**). Each bout lasted 5 s, with a 5 s interstimulus interval (ISI) during which the platform moved away by 1 cm and the object was out of reach of the test mouse. The total travel time for the platform was 1.2 s (for back and forwards). After this voluntary object touch session, the test mouse was returned to its cage to rest for at least a 60 min before being head-restrained again on the ball to undergo voluntary or forced social touch (randomized) session with a visitor mouse. A same-sex, same age (P60-90) novel WT mouse (for WT and *Fmr1*^*-/-*^ test mice) or a novel PBS mouse (for PBS and MIA test mice) was head-restrained inside the plexiglass tube on the stage. Following a 2 min period in the neutral position where the test mouse could see but not touch the stranger mouse, the motorized stage moved to the position for voluntary social touch (whisker-to-whisker) (**Movie 3**) or forced social touch (snout-to-snout) (**Movie 1**) for 5-20 bouts of each (also lasting 5 s with a 5 s ISI where the mouse on the platform moved out of reach of the test mouse). The test mouse was then returned to its cage for 24 h. On day #2, the mouse was placed back on the ball again for a 2 min baseline period followed by a 2 min period of no touch with a different stranger mouse. Depending on if the test mouse received voluntary or forced social touch on day 1, the mouse received 20 presentations of the alternate touch type with the second stranger mouse (**Fig. 1c**).

### QUANTIFICATION AND STATISTICAL ANALYSIS

#### Behavioral analyses

During the course of the assay, high-resolution videos (.mp4 files) were recorded of the test mouse’s eye, face, and body with 3 different cameras (Teledyne Flir, Blackfly S USB3) at 50 FPS for behavioral analyses. Avoidance running, aversive facial expressions, pupil diameter and locomotion were analyzed from these eye, face, and body videos (**Fig. 1d**). Running avoidance (backwards directed running), running speed and locomotion were analyzed from body videos using custom-written video analysis routines in MATLAB (**Fig. 2a, Fig. 2 Suppl. Fig. 1a**). Painted dots on the polystyrene ball (1 cm diameter) were used to measure the angle and distance based on the displacement of each dot at a frame to the closest dot 5 frames later (median angle and distance was calculated using all angles and distances for dots displaced for every 5 frames, or 0.1 s) (**Fig. 2a, Fig. 2 Suppl. Fig. 1a**). Pupil diameter was quantified using *Facemap* (Stringer et al., 2019) and MATLAB (**Fig. 3a, Fig. 3 Suppl. Fig 1a**). Aversive facial expressions (e.g., prolonged whisker protraction and orbital tightening) were also analyzed using *DeepLabCut*, as described (Mathis et al., 2018; Nath et al., 2019). Briefly, network was trained on images from the face videos to identify markers on the mouse’s whisker follicles. The displacement of the follicles was calculated using these markers to detect sustained (> 2 s) negative displacements from the resting position of the whiskers as aversive whisker movements **(Fig. 4a, Fig. 4 Suppl. Fig. 1a)**. We also quantified overall whisking during the assay by calculating the motion energy of whisking using *Facemap* (**Fig. 4 Suppl. Fig. 1b)**. To quantify orbital tightening or eye squinting, a neural network was trained on still images from videos of the face to reliably identify markers along the mouse’s eye. The area of the eye was calculated from these markers to quantify orbital tightening **(Fig. 5a, Fig. 5 Suppl. Fig. 1a)**.

#### Statistical analyses

Statistical tests were performed in Prism software (GraphPad). Statistical analyses of normality (Lilliefors and Shapiro Wilk tests) were performed on each data set; if data deviated from normality (p<0.05) or not (p>0.05), appropriate non-parametric and parametric tests were performed. For parametric two-group comparisons, a Student’s t-test (paired or unpaired) was used. For non-parametric tests, we used Mann-Whitney test (two groups) and the Kruskall-Wallis test (repeated measures). Multiple comparisons across touch conditions and genotypes/groups were analyzed using two-way ANOVA with post-hoc Bonferroni’s test. All experiments were conducted in at least two litters per genotype/group. Graphs either show data from each mouse per group or group means (averaged over different mice) superimposed on individual data points. In all figures, the error bars denote standard error of mean (s.e.m.).

## Abbreviations

AFE: aversive facial expression;
ASD: autism spectrum disorder;
MIA: maternal immune activation;
NDC: neurodevelopmental condition;
P: postnatal day.

## ACKNOWLEDGEMENTS

We are grateful to Gunvant Chaudhari for writing MATLAB scripts to detect ball motion, Will Zeiger and Nazim Kourdougli for help with the design of the behavioral apparatus. Kimberly Battista (https://www.battistaillustration.com) made the illustration in Fig. 1a. This work was supported by the following grants: R01NS117597 (NIH-NINDS), R01HD108370 and R01HD054453 (NIH-NICHD), Department of Defense (DOD, 13196175) awarded to C.P-C, Training in Neurotechnology Translation T32NS115753 (NIH), F31HD108043 (NIH/NICHD), and a graduate student fellowship from the Achievement Rewards for College Scientists Foundation to T.C., and the CARE Fellows Program to A.H.

## FIGURES LEGENDS

**Fig. 1, Suppl. Fig. 1:**
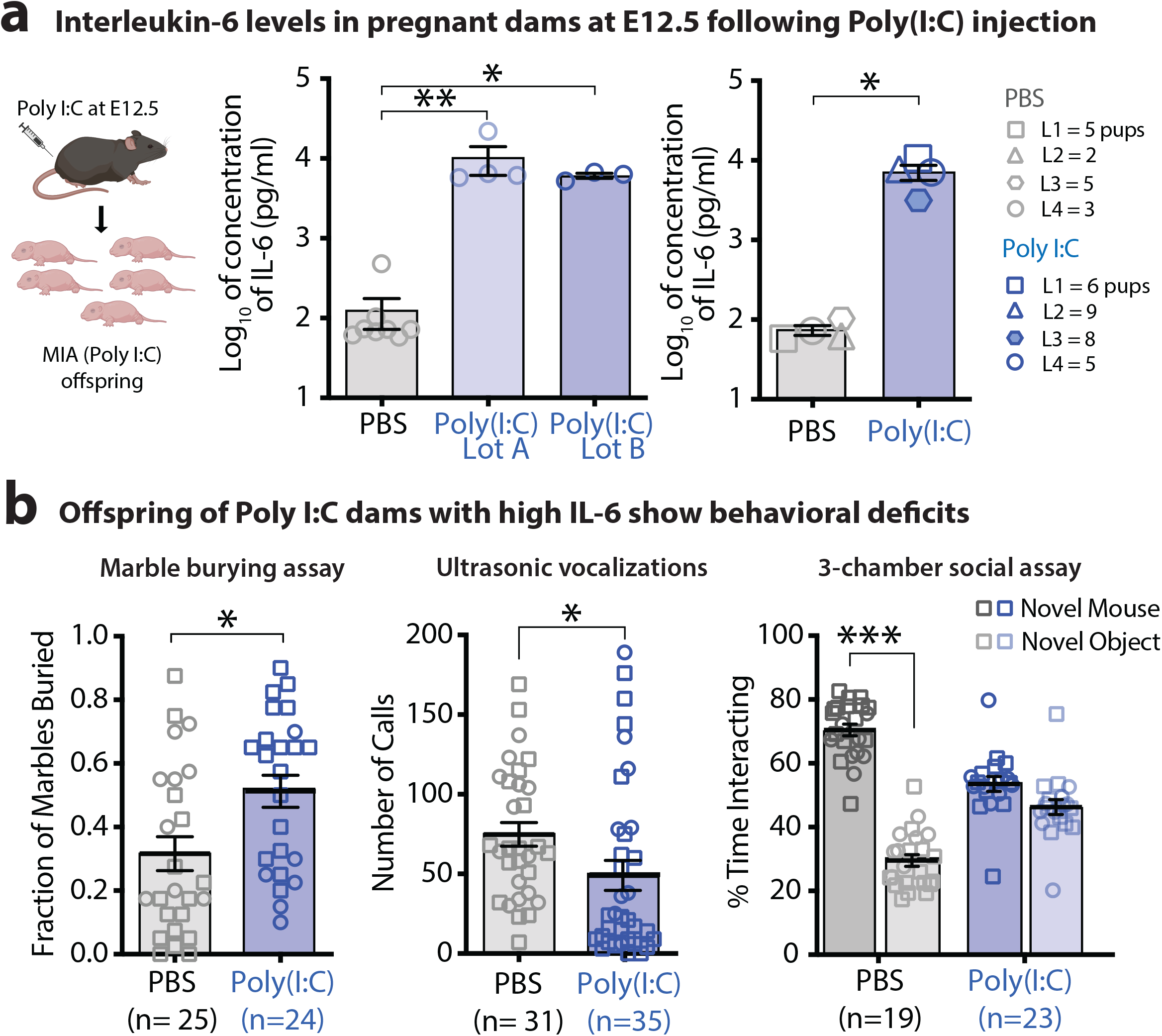
Interleukin-6 levels are higher in pregnant dams injected with Poly(I:C) and their offspring show expected behavioral deficits. **a**. MIA was induced in pregnant dams by intraperitoneally injecting Poly I:C at embryonic age 12.5 (E12.5). Interleukin-6 (IL-6) cytokine levels are higher in pregnant dams injected with Poly I:C at E12.5 compared to dams injected with PBS. Two different Poly I:C lots acquired from Sigma-Aldrich were tested and elicited significantly higher IL-6 levels in dams (left). These two lots were used to generate offspring for the behavioral testing in the social touch assay (right). ** p < 0.01, * p < 0.05 for Mann-Whitney test. **b**. Offspring of dams injected with Poly I:C from Lot A and B showed increased fraction of marbles buried in the marble burying assay at P60-90, reduced ultrasonic vocalizations recorded at P7-9, and no difference in preference for a novel mouse versus novel object in the 3-chamber social interaction assay. Squares = males, circles = females in panel b. *** p < 0.001, *p < 0.05, unpaired t-test for marble burying assay and USVs, two-way ANOVA with Bonferroni’s for 3-chamber social assay.

**Fig 2. Suppl. Fig. 1:**
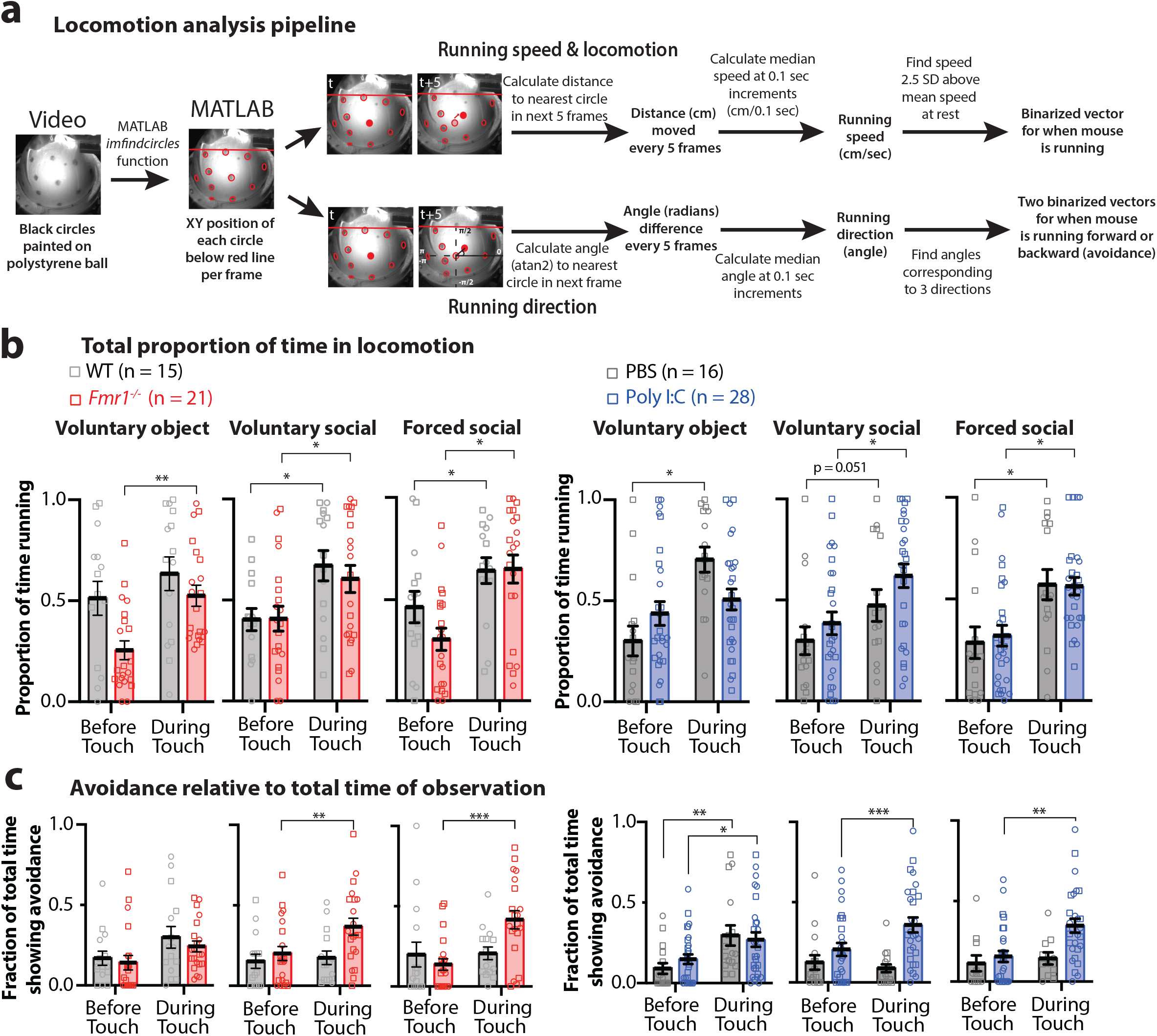
Locomotion & avoidance increase in response to social & object touch. **a**. Detailed analysis for locomotion, running speed and direction. Speed (cm/s) is calculated using the distance moved of a circle at time t to the closest circle in pixel space in time t+5 every 5 frames (0.1 s). Direction of circle movement at time t to t+5 frames is calculated from the angle between the circle at time t as the origin point relative closest circle at t+5 in pixel space. Median speed and angle is calculated from speeds and angle displacements of all circles every 5 frames (0.1 s). Locomotion calculated by finding running speeds 2.5 standard deviations above mean speed at rest. Example circle (red filled in time t and pink filled in t+5) moves to the right and up (red filled in time t+5, leftwards avoidance). Circles above red line are excluded from detection. **b**. Locomotion increased during social touch compared to before touch but doesn’t differ between groups. **c** Proportion of running avoidance (backwards to left or right) over total time of social touch is significantly higher during social touch in Fmr1-/- and MIA mice. Squares = males, circles = females. *** p <0.001, ** p < 0.01, *p < 0.05, two-way ANOVA with Bonferroni’s.

**Fig 2. Suppl. Fig. 2:**
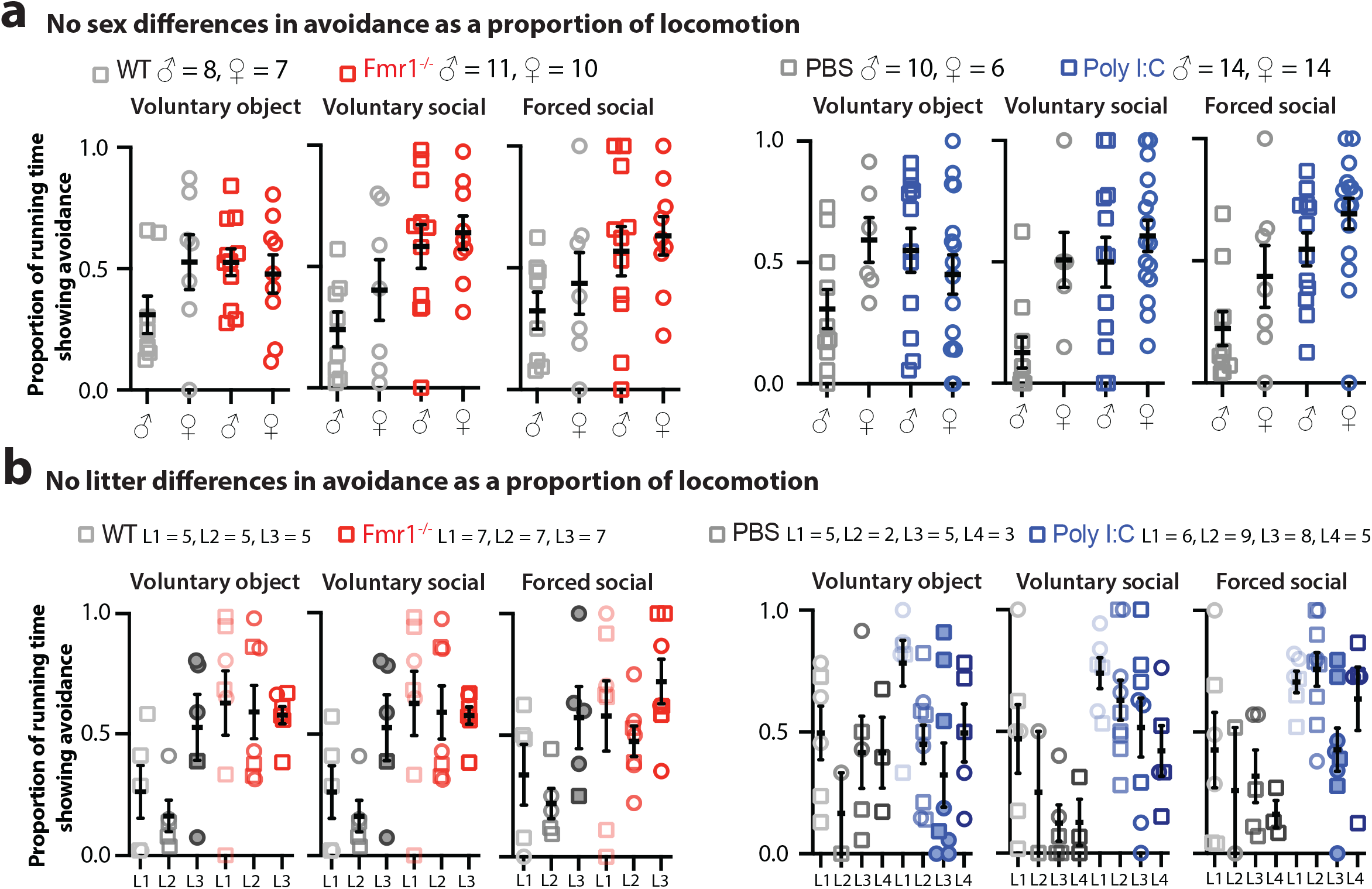
Running avoidance does not differ based on sex across groups and does not differ across litters in each group during object and social touch. **a**. Running avoidance as proportion of total running was not significantly higher in females compared to males for across all groups. **b**. No difference in running avoidance across litters in each group during object and social touch. Statistical analysis was performed on the sex differences only. One litter (L3) for WT and MIA (filled circle) deviate in their responses relative to litters in the same group. Squares = males, circles = females. p > 0.05 for Kruskal-Wallis test. Vol = voluntary. L1-4 denotes litters.

**Fig 3. Suppl. Fig. 1:**
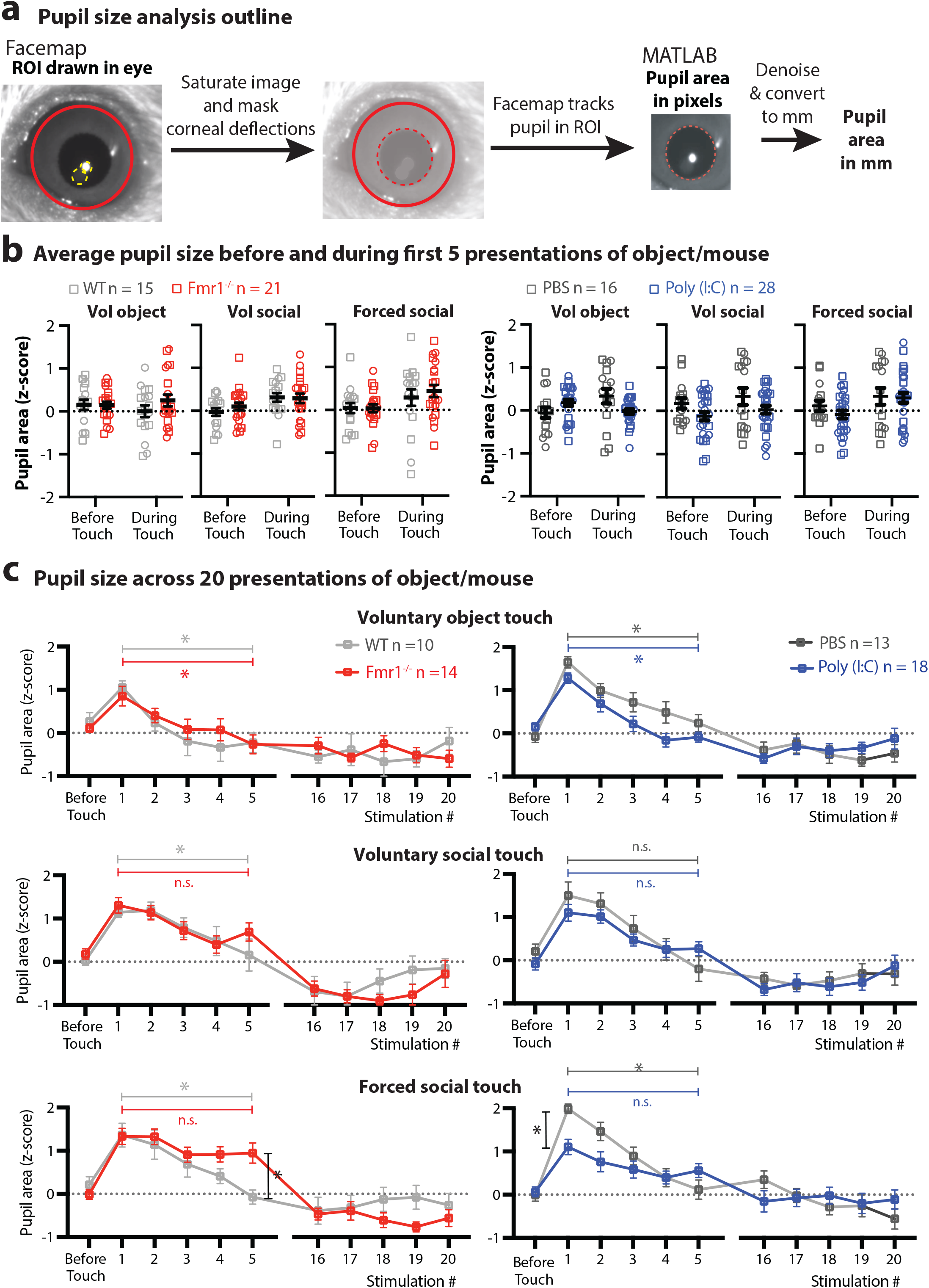
Pupil size does not differ before touch but pupil size decreases more slowly after repeated socials presentations in ASD mice. **a**. Analysis for pupil size using Facemap. A region of interest (ROI) is drawn in the Facemap graphical user interface (GUI) in Python (red circle). Circles are also drawn on corneal deflections (yellow dashed circle), which are then masked by Facemap to enhance pupil detection. Facemap then detects the pupil within the ROI (red dashed circle) and generates pupil area in pixels.In MATLAB, the pupil area signal is denoised using MATLAB’s smooth function and converted to mm based on ratio of pixels to mm in the camera’s depth of field. **b**. Pupil size doesn’t differ between groups before social touch and as an average of all 5 stimulations of social touch (during social touch) based on post-hoc Bonferroni’s. **c**. A subset of mice were tested for up to 20 presentations. In this subset, Fmr1-/-mice show significantly larger pupil dilation at the fifth stimulation but decrease to WT levels by the 16th stimulation. MIA mice in this subset do not increase pupil size at the onset of the first stimulation at the onset of the first forced social touch as PBS mice but show less pupil constriction when comparing the 1st to the 5th stimulation. There were no differences in pupil dilation across times between groups for voluntary object and social touch and all groups adapt by the 16^th^ stimulation to pupil size before touch. Squares = males, circles = females. *** p < 0.001, ** p < 0.01, *p < 0.05, two-way ANOVA with Bonferroni’s. Vol = voluntary.

**Fig. 3 Suppl. Fig. 2:**
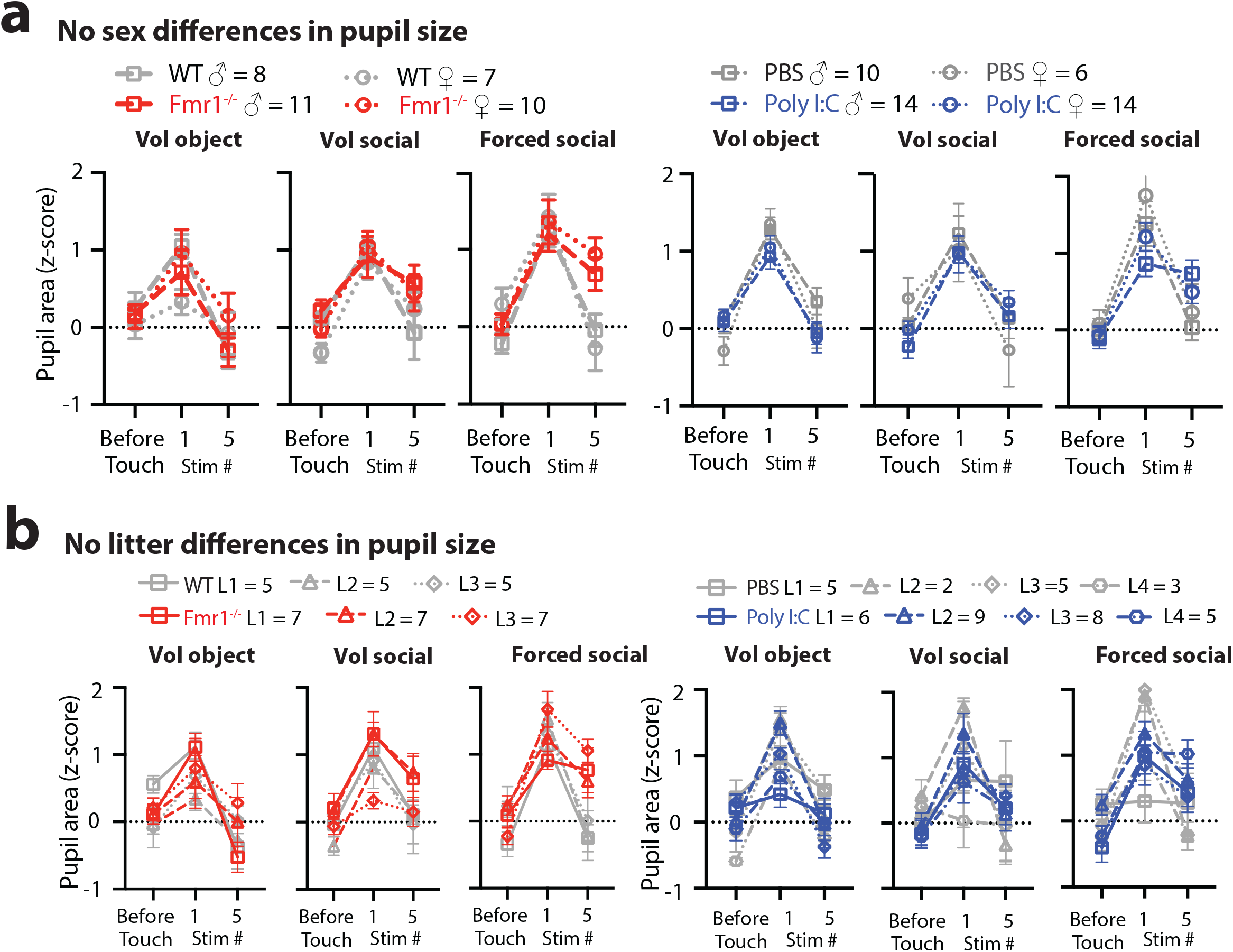
No sex or litter differences in pupil size across groups. **a**. Change in pupil dilation across time (before touch versus 1st and 5 stim of touch) was not different between males and females in each group. **b**. Pupil dilation changes did not differ across litters in each group during object and social touch. Statistical analysis was performed on the sex differences only. p > 0.05 for Kruskal-Wallis test. L1-4 denotes litters. Vol = voluntary.

**Fig. 4 Suppl. Fig. 1:**
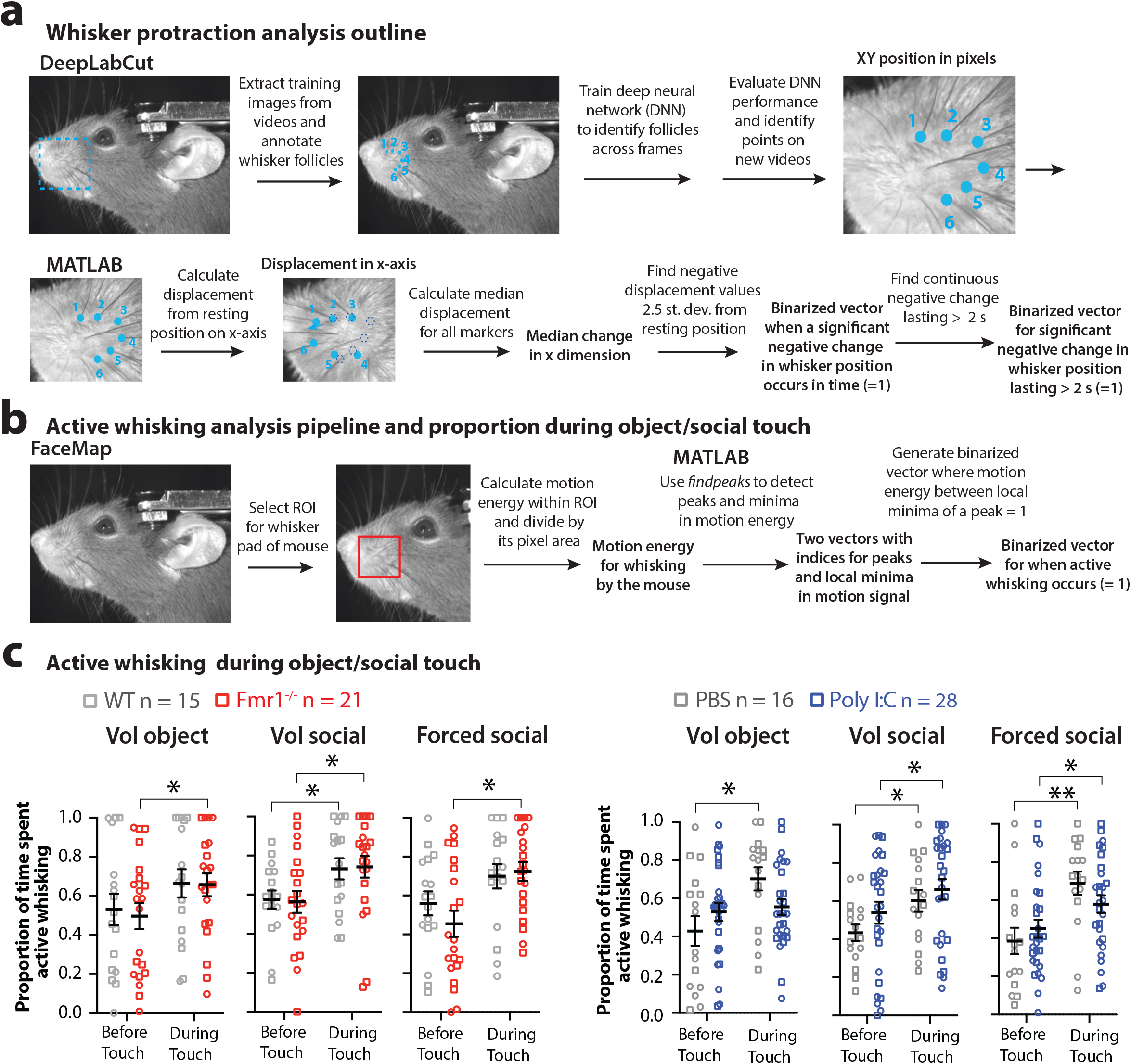
Analysis of whisker protraction and active whisking; No differences in active whisking in mouse models of autism. **a**. Whisking protraction was determined by training a deep neural network (NN) in DeepLabCut to detect 6 whisker follicles from a set of training video frames (randomly chosen frames). We indicated the 6 follicles in the training image using DeepLabCut. After training the NN and evaluating its performance, we processed full videos, which generated the x-y position of each whisker follicle in pixel space. We then calculated the change of each follicle position relative to its resting position along the x-axis. Negative changes in follicle position over time that were 2.5 standard deviations above mean resting position and lasted more than 2 seconds were denoted as periods of aversive whisker protraction. **b**. To quantify periods of active whisking we used FaceMap. We denoted the whisker pad as a region of interest (ROI) and tracked changes in pixel value within the ROI as the motion energy for whisking. We then used MATLAB’s findpeaks function to identify peaks and local minima in the motion energy signal. We identified periods of active whisking as timepoints that were between the local minima of a peak. **c**. Active whisking did not differ between Fmr1-/- and MIA mice and their controls both before and during object and social touch. There was, however, a significant increase in mean whisking during the 5 stimulations compared to before touch. Squares = males, circles = females. * p < 0.05, ** p < 0.01 for two-way ANOVA with Bonferroni’s. Vol = voluntary.

**Fig. 4 Suppl. Fig. 2:**
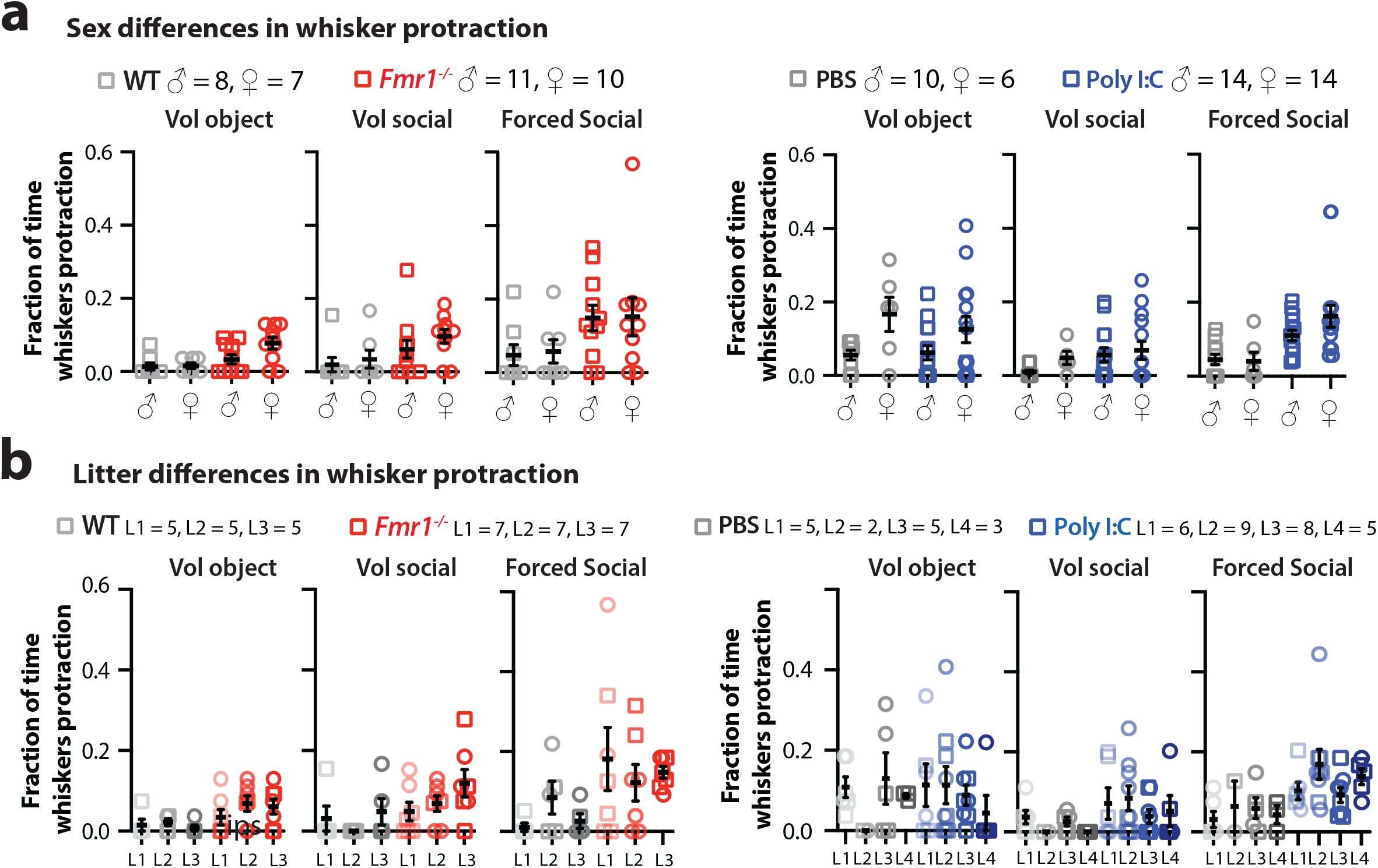
No sex or litter differences for aversive whisking in social touch assay. **a** Proportion of time spend aversive whisking was not significantly different between males and females in each group. **b** Aversive whisking did not differ across litters in each group during object and social touch. Statistical analysis was performed on the sex differences only. Squares = males, circles = females. p > 0.05 for Kruskal-Wallis test. Vol = voluntary. L1-4 denotes litters.

**Fig. 5: Suppl. Fig. 1:**
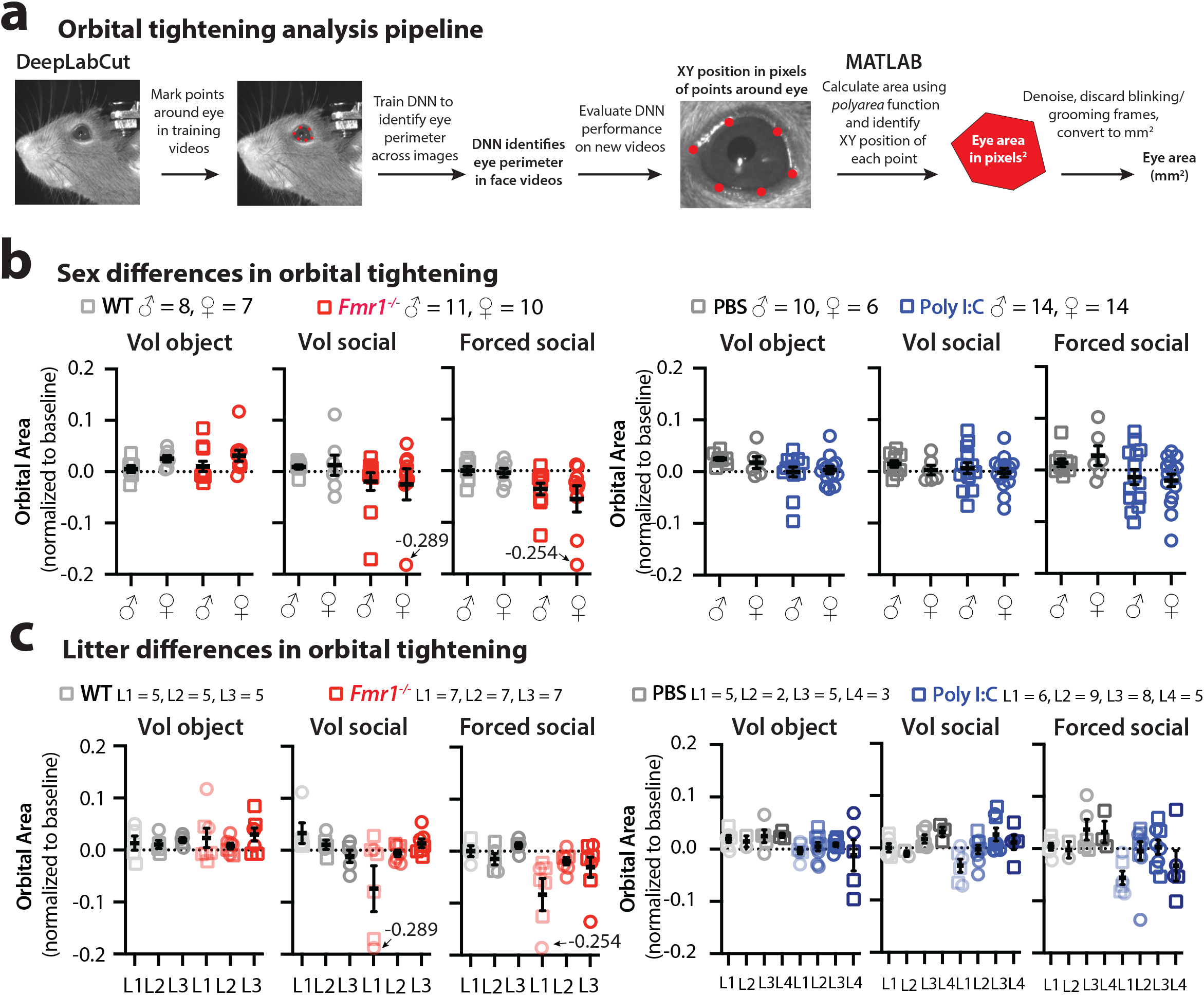
No sex or litter differences for orbital tightening. **a**. Orbital tightening is determined by training a deep neural network (NN) in DeepLabCut to detect 6 points along the eye from a set of training images (frames randomly chosen from videos). In DeepLabCut, we indicated the 6 eye points in the training image using DeepLabCut’s graphical user interface (GUI). After training the NN and evaluating its performance, we inputed videos into the Deep NN, which outputs the XY position of each point on the eye in pixel space. We then used MATLAB to generate a polygon connecting the six dots and calculated the area of that polygon as the orbital area. Orbital area in pixels was converted to mm based on ratio of pixels to mm in the camera’s depth of field and then normalized to the orbital area during baseline (no object or mouse is present). **b**. Orbital area was not significantly different between males and females in each group **c** Orbital tightening did not differ across litters in each group during object and social touch. Statistical analysis was performed on the sex differences only. p > 0.05 for Kruskal-Wallis test. Vol = voluntary. L1-4 denotes litters.

**Movie 1: Five stimulations by forced social touch**

**Movie 2: A single stimulation of voluntary object touch**

**Movie 3: A single stimulation of voluntary social touch**

**Movie 4: Pupil dilation in a mouse during social touch**

**Movie 5: Aversive facial expressions in a mouse during social touch**

